# Boosting of SARS-CoV-2 immunity in nonhuman primates using an oral rhabdoviral vaccine

**DOI:** 10.1101/2021.10.16.464660

**Authors:** Kah-Whye Peng, Timothy Carey, Patrycja Lech, Rianna Vandergaast, Miguel Á. Muñoz-Alía, Nandakumar Packiriswamy, Clement Gnanadurai, Karina Krotova, Mulu Tesfay, Christopher Ziegler, Michelle Haselton, Kara Sevola, Chase Lathrum, Samantha Reiter, Riya Narjari, Baskar Balakrishnan, Lukkana Suksanpaisan, Toshie Sakuma, Jordan Recker, Lianwen Zhang, Scott Waniger, Luke Russell, Christopher D. Petro, Christos A. Kyratsous, Alina Baum, Jody L. Janecek, Rachael M. Lee, Sabarinathan Ramachandran, Melanie L. Graham, Stephen J. Russell

## Abstract

An orally active vaccine capable of boosting SARS-CoV-2 immune responses in previously infected or vaccinated individuals would help efforts to achieve and sustain herd immunity. Unlike mRNA-loaded lipid nanoparticles and recombinant replication-defective adenoviruses, replicating vesicular stomatitis viruses with SARS-CoV-2 spike glycoproteins (VSV-SARS2) were poorly immunogenic after intramuscular administration in clinical trials. Here, by G protein trans-complementation, we generated VSV-SARS2(+G) virions with expanded target cell tropism. Compared to parental VSV-SARS2, G-supplemented viruses were orally active in virus-naive and vaccine-primed cynomolgus macaques, powerfully boosting SARS-CoV-2 neutralizing antibody titers. Clinical testing of this oral VSV-SARS2(+G) vaccine is planned.

## Introduction

Control of the SARS-CoV-2 pandemic will depend in part upon the global deployment of effective vaccines [1]. However, all currently approved SARS-CoV-2 vaccines are administered via intramuscular injection which increases the cost and complexity of vaccination programs, contributes to vaccine hesitancy and fails to stimulate mucosal immunity [2, 3]. Also, protective antibody titers are known to wane after successful vaccination (or after natural infection), such that periodic booster immunizations will be needed to sustain immunity, particularly in vulnerable individuals, so long as the virus remains endemic in the human population [4–6].

Since oral and nasal mucosal surfaces are the primary portal of entry for SARS-like coronaviruses, the induction of mucosal immunity (via secretory IgA) may be the best way to prevent the virus from ever getting a foothold in the upper airways, thereby eliminating the risk of asymptomatic infection and virus shedding in vaccinated individuals which could aid virus transmission [7]. Available data from previous studies suggests that mucosal protection can be achieved most readily via mucosal vaccination using an oral or intranasal delivery route [3] which has been exemplified in previously licensed vaccines (e.g. Rotarix^®^ and oral polio vaccines). From a cost, convenience and simplicity perspective, the oral route is preferred over the intranasal route since it does not require the use of a specialized delivery device (e.g. nebulizer). Also, from a safety perspective oral delivery avoids the risk that a live viral vaccine administered intranasally might spread into the brain via the olfactory neurons which pass through the cribriform plate at the apex of the nasal cavity [8]. Finally, compared to available injectable vaccines, oral vaccines are much preferred by children and adolescents whose high prevalence of vaccine hesitancy is linked to fear of needles [2, 9].

Here, to develop an orally active viral vaccine capable of expressing the immunogenic SARS-CoV-2 spike glycoprotein in oral mucosa, we exploited the known tropism of vesicular stomatitis virus (VSV). Vesicular stomatitis is a self-limited illness of cattle and other ungulate species characterized by blistering of the oral mucosa and hooves [10]. The disease is endemic in Central America and the Southern United States, and the causative agent is vesicular stomatitis virus (VSV), a mosquito borne Vesiculovirus belonging to the family Rhabdoviridae. Human exposures to VSV infected cattle lead to asymptomatic seroconversion or sometimes a brief febrile illness with associated oral blistering [10]. VSV seroprevalence is very low in the human population, even in endemic areas [11]. VSV is a bullet-shaped virus with a lipid envelope and a single stranded negative sense RNA genome of approximately 11kb, comprising five genes encoding nucleocapsid (N), phosphoprotein (P), matrix (M), glycoprotein (G) and polymerase (L) proteins. Virus entry is mediated via the interaction of the viral G protein with the low density lipoprotein (LDL) receptor, ubiquitously expressed on mammalian cells [12].

To generate an orally active VSV-SARS2(+G) vaccine we substituted the G cistron of an Indiana strain VSV genome with a codon-optimized sequence encoding the Wuhan strain SARS-CoV-2 spike glycoprotein (S) and amplified the recombinant virus on Vero cells transiently expressing the G protein (Fig. 1A). To facilitate S protein incorporation and G-independent propagation of the recombinant VSV-SARS2 viruses, a 19 amino acid S protein cytoplasmic tail truncation was introduced into the S protein [13]. The virus rescued from this construct was able to propagate autonomously, albeit slowly, with characteristic syncytial cytopathic effect on A549 cells transduced with the ACE2 receptor, but not on parental A549 cells (Fig. 1B). Autonomously replicating non-G-deleted VSVs encoding the SARS-CoV2 spike protein that were generated in parallel were found to replicate efficiently on ACE2 negative cells, but rapidly lost expression of the spike protein, primarily via point mutations in key upstream regulatory sequences (data not shown). The stably and autonomously replicating VSV-SARS2 virions were next propagated on ACE2 receptor positive Vero cells transfected with a VSV-G expression plasmid to trans-pseudotype them with the G protein, which is also known to drive more efficient virus budding [14]. G-protein incorporation led to an approximately 10-fold enhancement of the infectious titer of the VSV-SARS2(+G) virus preparations (titers of 2 ×10^9^ TCID_50_ per ml were achieved for the G supplemented viruses versus maximum titer of 5.1 × 10^7^ TCID_50_ without G protein. In a parallel study, incorporating the SARS-CoV-2 spike coding sequences as an additional cistron (that is, without deletion of G) proved to be highly unstable since their replication was not dependent on the integrity of the SARS-CoV-2 spike.

**Figure 1:**
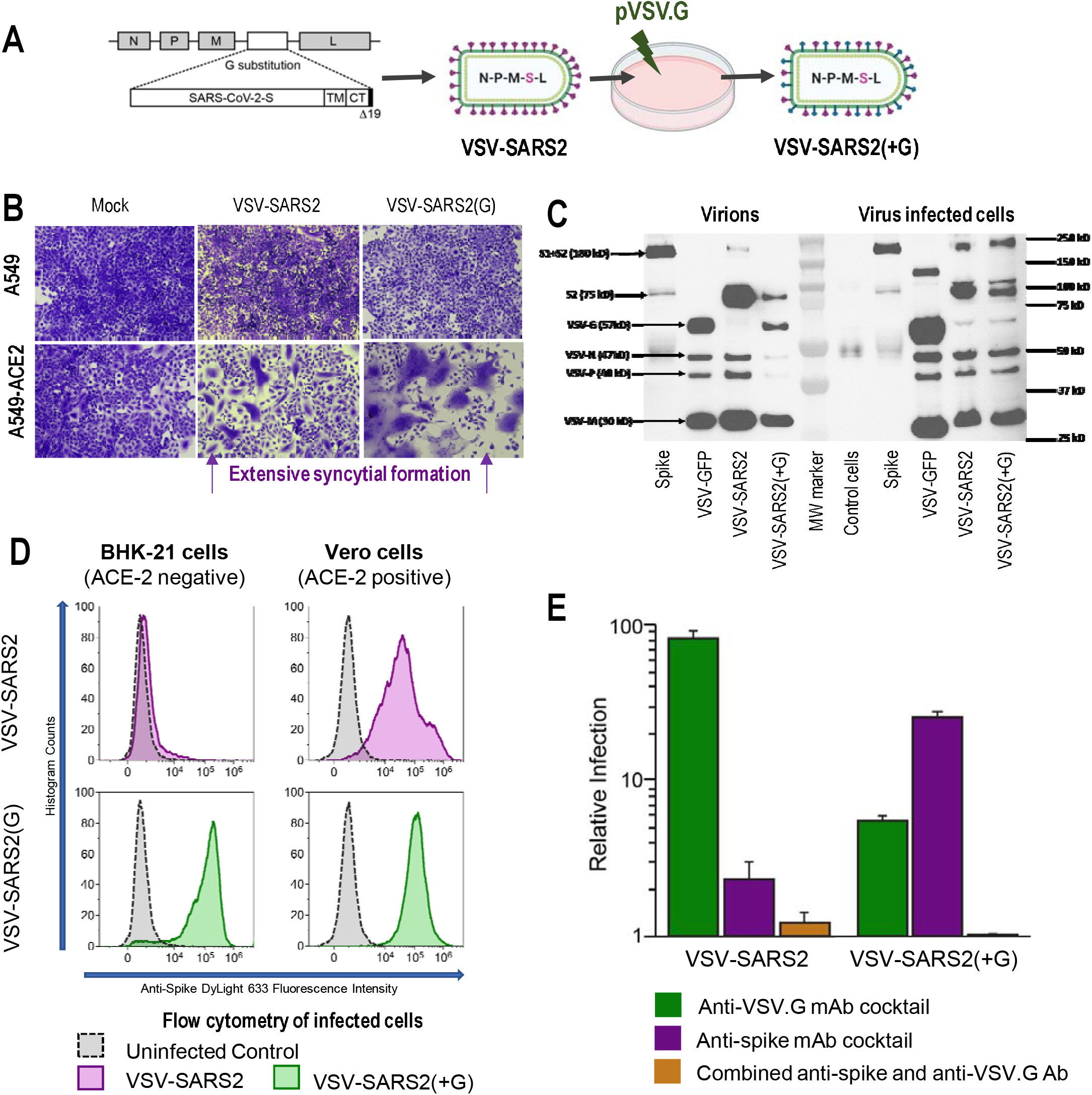
Generation and characterization of VSV-SARS2 and VSV-SARS2(+G) viruses. (**A**) Schematic of the VSV-SARS2 genome and nomenclature of viruses. (**B**) Crystal violet stained A549 cells and A549-ACE2 cells at 24h after virus infection. Note the extensive syncytial formation in the VSV-SARS2 and VSV-SARS2(G) infected A549-ACE2 expressing cells. (C) Immunoblot of virions prepared in untransfected or G-plasmid transfected Vero cells, and of lysates from corresponding virus infected cells were probed with polyclonal VSV antiserum and a monoclonal antibody against the S2 domain of the SARS-CoV-2 spike glycoprotein. The S2 shadow band in the cell lysates may be a glycosylation variant. (**D**) Flow cytometry analysis of anti-spike stained virus infected cells. (**E**) Neutralization of the infectivity VSV-SARS2 (+/- G) by a cocktail of anti-G and/or anti-S monoclonal antibodies.

Biochemical characterization of the recombinant virus particles confirmed the incorporation a partially cleaved S protein in pure VSV-SARS2 virions, and the co-incorporation of S and G proteins in the G-pseudotyped VSV-SARS2(+G) virions (Fig. 1C). SARS-CoV2 S protein expression after infection with VSV-SARS2 or VSV-SARS2(+G) was confirmed by western blotting and/or flow cytometry of ACE2 negative baby hamster kidney cells (BHK) and ACE2 positive Vero cells after infection by the respective viruses (Fig. 1C and D). VSV-SARS2 virions were neutralized by a cocktail of monoclonal anti-S SARS-CoV-2 neutralizing antibodies, but not by anti-G antibodies, indicating that they enter cells exclusively via their displayed S protein (Fig. 1E). In contrast, after treatment with anti-S or anti-G antibodies, G-pseudotyped VSV-SARS2(+G) virions retained 20% or 4% infectivity respectively but were more fully neutralized when the anti-S and anti-G antibodies were used in combination (Fig. 1E).

To facilitate in vivo studies in non-human primates (NHP), and possible clinical translation, mid-scale preparations of the VSV-SARS2 and VSV-SARS2(+G) viruses were generated using adherent GMP-qualified Vero cells as the substrate. Crude viral supernatants from these production runs were clarified by centrifugation and filtration, treated with benzonase, and purified by tangential flow filtration prior to cryopreservation. Viral stocks were titrated post-thawing by TCID_50_ assay on Vero cell monolayers, then characterized for S and G protein incorporation and endotoxin content.

Expression of ACE2 receptors is not prominent on primate myocytes [14]. However, based on a published report demonstrating protective immunity to SARS-CoV-2 in hamsters vaccinated by intramuscular administration of a G-deficient VSV-ΔG-SARS-CoV-2 construct [15], we tested our own G-deficient VSV-SARS2 construct by intramuscular (IM, 10^7^ TCID_50_) vaccination of six virus-naive cynomolgus macaques (Fig. 2A). Compared to unvaccinated control animals, all monkeys in this experimental group developed detectable anti-S antibody responses by 4 weeks after vaccination (Fig. 2, B, C). However, the absolute magnitude of the anti-spike IgG and neutralizing antibody titers observed in these animals were very low relative to mean human convalescent titers (Farcet. In parallel with this intramuscular study, 10 mls of the virus (10^7^ TCID_50_) was administered into the oral cavity of animals (PO) under brief sedation. Immune responses to the oral application of the virus were not detected in any of the animals in this group (Fig. 2, B, C). The conclusion of this preliminary study in cynomolgus macaques was that the G-deficient VSV-SARS2 virus was weakly immunogenic when administered by the intramuscular route and is minimally immunogenic by the oral route.

**Figure 2:**
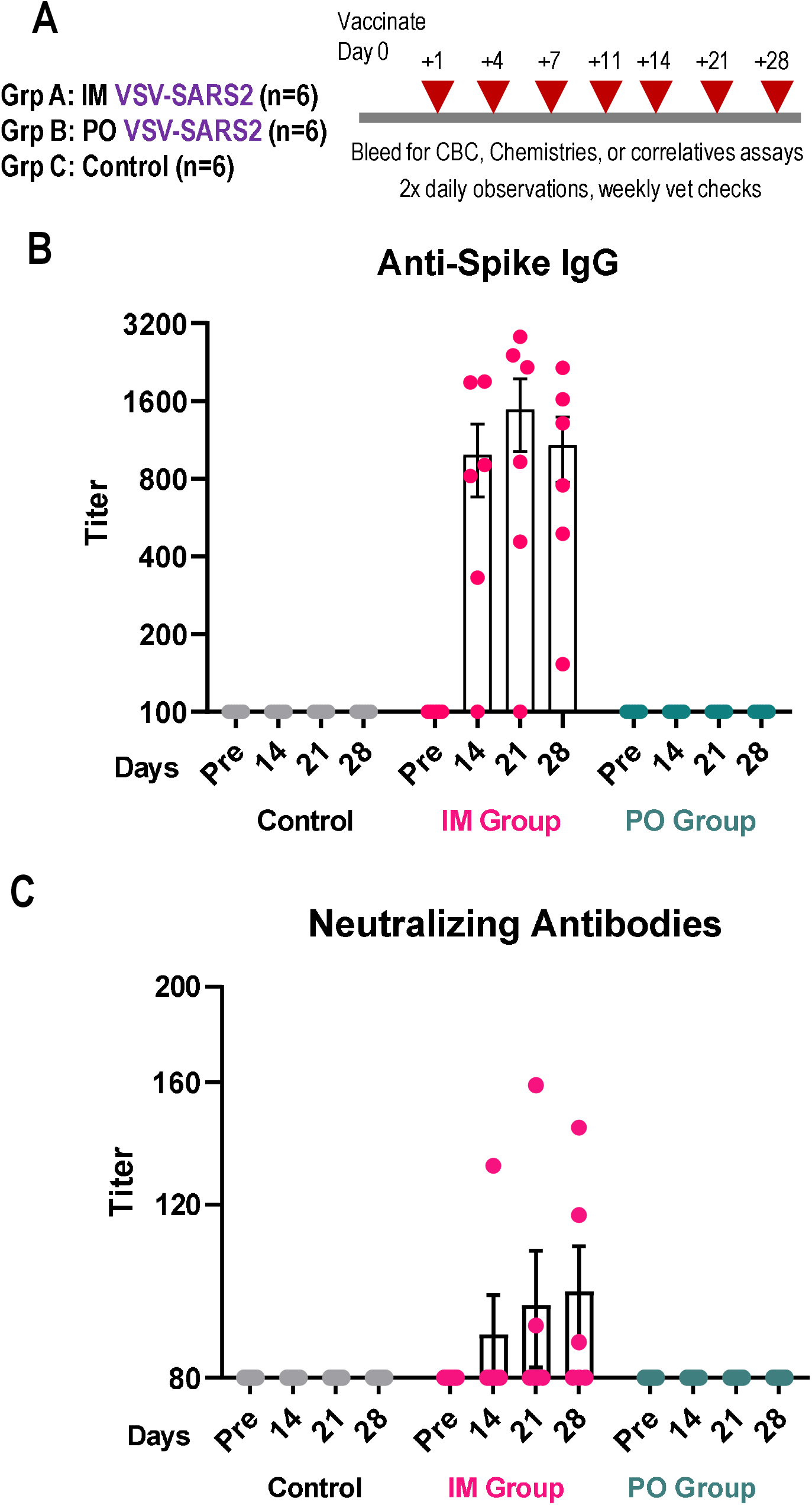
VSV-SARS2 is poorly immunogenic in cynomolgus macaques (IM and oral routes). (**A**). Experimental design showing groups of six cynomolgus macaques vaccinated by intramuscular (IM) or oral (PO) routes using 10^7^ TCID_50_ VSV-SARS2 virus. Another six animals received vehicle control via both IM and PO routes. Baseline and post-vaccination samples were harvested at indicated timepoints. (**B**) IgG anti-spike antibody titers by ELISA. (**C**) Neutralizing antibody titers using a live SARS-CoV-2 spike VSV pseudovirus (IMMUNO-COV™) assay [17]. Time points are before (pre) and at 2, 3, and 4 weeks post vaccination. A log2 scale is used for the y axis.

To determine whether G protein supplementation could enhance the potency of the recombinant VSV vaccine, we administered VSV-SARS2(+G) to a total of four virus-naive cynomolgus macaques, either by IM injection (n=2) or PO (n=2). Both IM vaccinated animals and one of the orally vaccinated animals rapidly generated high IgG and neutralizing antibody titers against the SARS-CoV-2 spike glycoprotein (Fig. 3, A, B). Neutralizing antibody titers were well maintained with no significant decline in two of the three responding animals until the end of the study (six months post vaccination) indicating excellent durability of the immune response to this vaccine formulation. Interestingly, even though the VSV-SARS2(+G) virus does not encode the G protein, both IM-vaccinated animals generated low titers of G-reactive VSV neutralizing antibodies (Fig. 3C), indicating that even the small amount of G protein carried by the injected virus particles was sufficient to induce an immune response via this route. In contrast, anti-G antibodies were not detected in the orally vaccinated animals (Fig. 3C), indicating that particle-associated G protein may be less likely to provoke an immune response when the viruses are administered orally.

**Figure 3:**
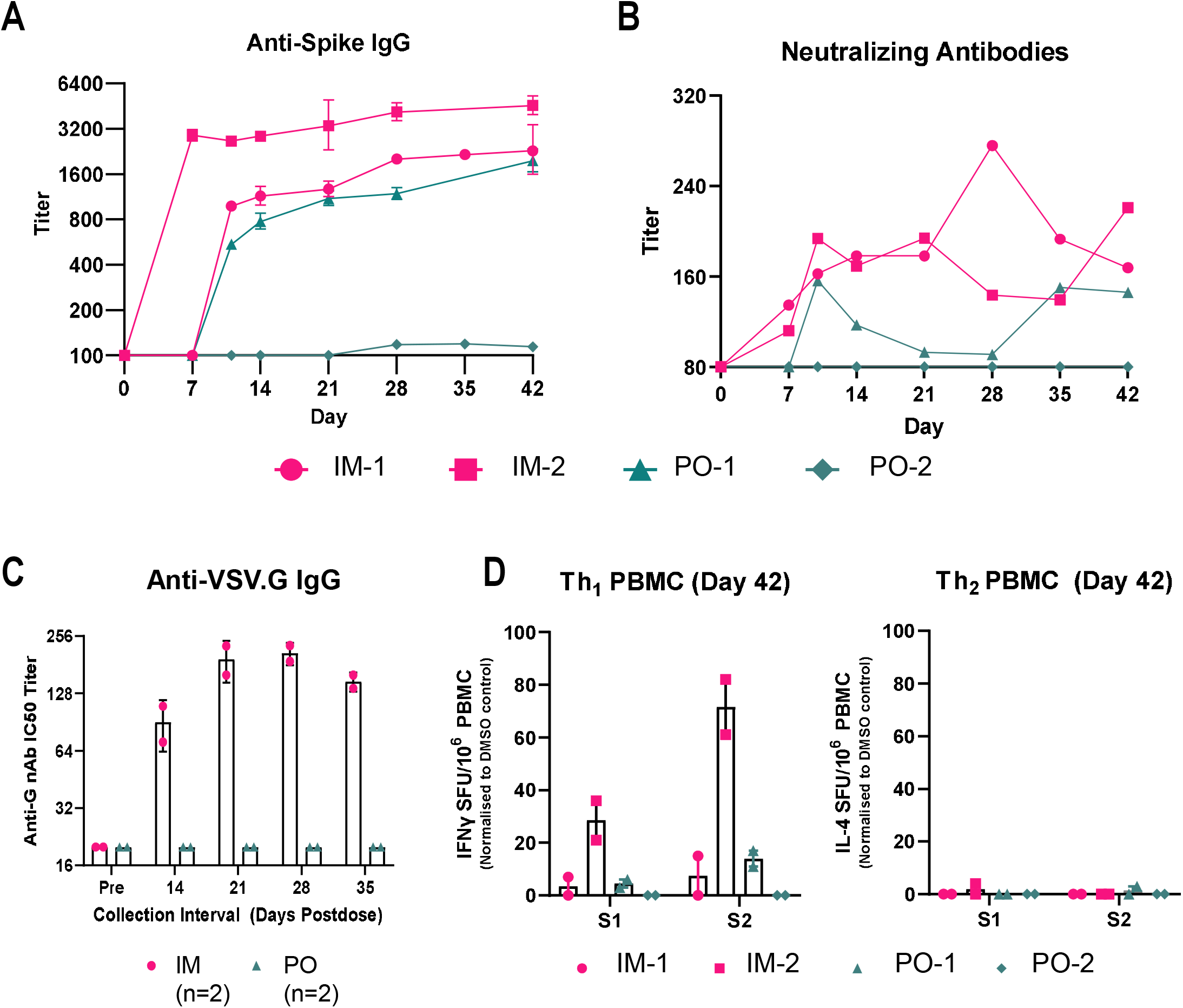
VSV-SARS2(+G) is powerfully immunogenic in cynomolgus macaques. 4 NHP were given the VSV-SARS(+G) vaccine by IM (IM-1, IM-2) or PO routes (PO-1, PO-2). Titers of anti-spike (A) IgG and (B) neutralizing antibodies against a live SARS-CoV-2 spike pseudovirus were measured at various time points. (**C**) Anti-VSV.G IgG titers in the orally or IM vaccinated animals. (**D**) Th1 (IFN-gamma secreting) and Th2 (IL-4 secreting) ELISPOT assay for spike antigen reactive T cells in peripheral blood mononuclear cells (PBMC) at day 42.

Analysis of T cell responses to the SARS-CoV-2 S protein in vaccinated macaques that seroconverted demonstrated an increase in spike antigen specific interferon gamma (IFN-γ) ELISPOT positive T cells but not IL4 positive T cells, indicating Th_1_ skewing (Fig. 3D). In light of the importance of this observation, we conducted confirmatory studies in immunized immune competent mice by intraperitoneal injection of the VSV-SARS2(+G) virus. As shown in supplemental figure 1, all vaccinated mice had a robust anti-spike antibody response with predominance of IgG2a antibodies compared to IgG1, further confirming the potent immunogenicity of the platform with associated Th_1_ skewing [16] of the immune response in the mouse model.

To determine whether the orally administered VSV-SARS2(+G) vaccine might have the ability to refresh and amplify the SARS-CoV-2 immune responses of individuals previously infected or vaccinated, but with low or waning titers of SARS-CoV-2 neutralizing antibodies, we orally boosted all six cynomolgus macaques from Experiment 1, Group A, 42 days after they received the intramuscular G-deficient VSV-SARS2 vaccine (Fig. 4A). Four animals received a high dose (10^9^ TCID_50_) of the oral vaccine and two animals received a low dose (5×10^6^ TCID_50_). Logarithmic increases in the total IgG and neutralizing antibody titers were observed in all four animals in the high dose group within a week of receiving the oral boost (Fig. 4, B-D). Similar boosting of the antibody response was observed in one of the two animals in the low dose group (Fig. 4, B-D). Neutralizing antibody titers were concordant (fig. S2) between the vesiculovirus (Fig. 4C) and lentivirus pseudovirus assays (Fig. 4D). In addition, we have previously shown that the neutralizing antibody assay using the vesiculo-pseudovirus has good correlation (R=0.89, p < 0.0001) with the classical plaque reduction neutralizing titer PRNT assay using the BSL3 live SARS-CoV-2 virus [17]. Analysis of peripheral blood mononuclear cells revealed significant increases in S protein specific IFNγ-secreting T cells in these orally boosted animals, but not of IL4-secreting T cells in a ELISPOT assay (Fig. 4E). Significantly, anti-G antibodies were not detected in any of the orally boosted animals (Fig. 4F), suggesting that it may be possible to use the G-pseudotyped platform repeatedly in the same individual to boost immunity against diverse coronavirus spike glycoproteins. Analysis of buccal and bronchoalveolar lavage (BAL) fluid obtained at euthanasia from these orally boosted and control animals revealed a substantial increase in the level of spike specific IgA as an indicator of mucosal immunity (Fig. 4, G and H).

**Figure 4:**
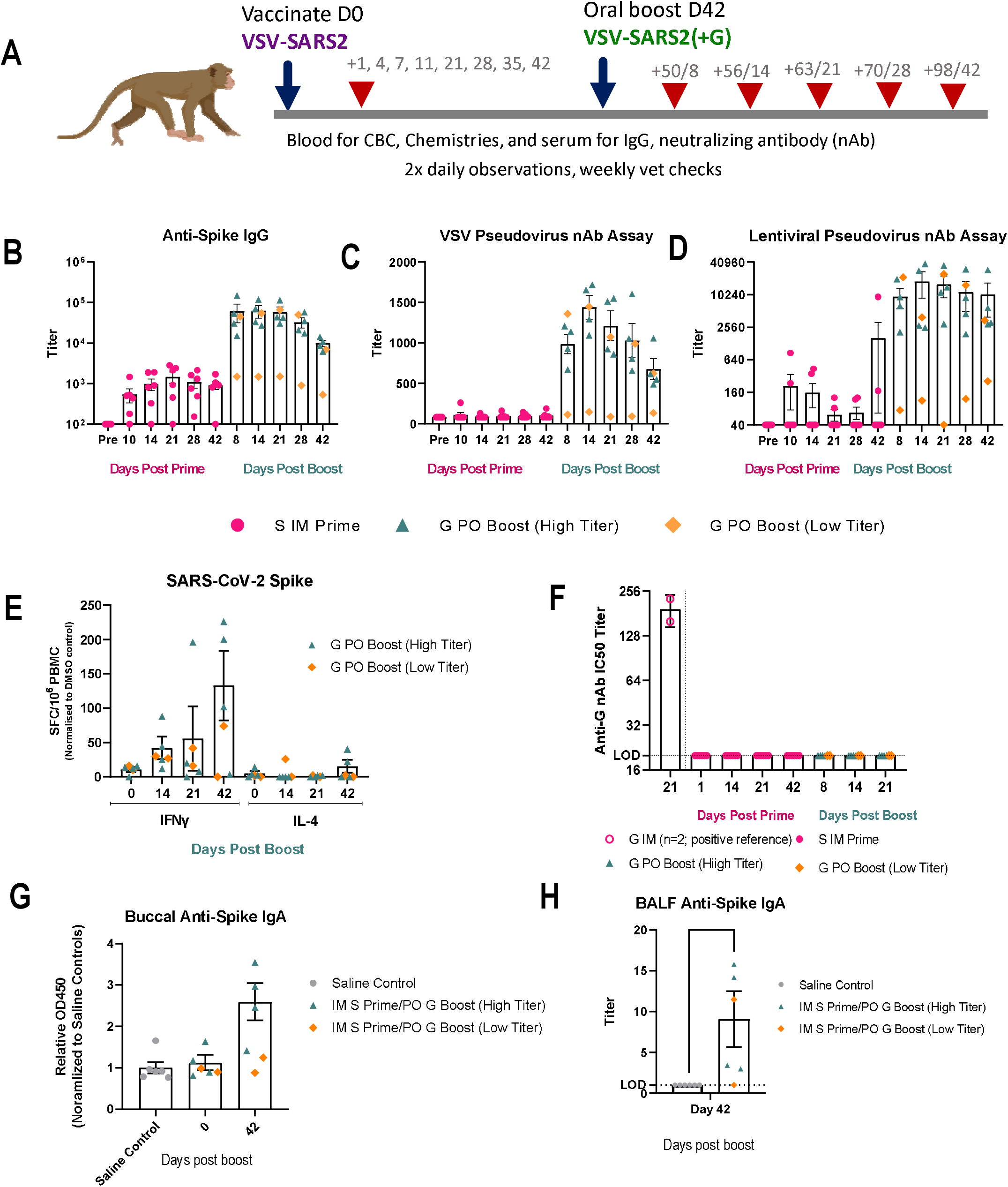
Oral VSV-SARS2(+G) powerfully boosts the immune response in vaccine-primed cynomolgus macaques. (**A**) Experimental design. Animals received initial VSV-SARS2 vaccine by IM or PO routes, and at day 42, the IM vaccinated NHP received an oral boost of VSV-SARS2(+G). 4 NHP received the high dose of 10^9^ TCID_50_ and 2 NHP received the lower dose of 5×10^6^ TCID_50_ VSV-SARS2(+G) (**B**) IgG titers against spike. (**C**) Neutralizing antibody titers using the live SARS-CoV-2 spike VSV pseudovirus and (**D**) a lentiviral pseudovector displaying the spike protein. (**E**) IFN-gamma and IL-4 ELISPOT assay for spike antigen reactive T cells from peripheral blood mononuclear cells (PBMC) after oral boost with VSV-SARS2(+G). G= VSV-SARS(+G) virus, S=VSV-SARS2 virus (**F**) Anti-VSV.G IgG antibody titers in boosted animals. (**G**) Anti-spike IgA titers in bronchoalveolar lavage fluid (BALF) and (**H**) buccal fluid of animals.

The tolerability of VSV-SARS2 and VSV-SARS2(+G) vaccines was examined in all NHPs via physical examination, behavioral observations, complete blood counts, and serum chemistries, with a complete necropsy at the end of the study. Body weights were stable with a gain trend and there were no significant changes in the serum C-reactive protein, an acute inflammation marker (fig. S3). Mild increases in body temperature were observed within the normal range for vaccinated animals and were observed with equivalent frequency in sham-vaccinated animals. A low incidence of vomiting and inappetence was observed following sedation in both sham controls and vaccinated animals which is a well-known side effect of the drugs used for sedation. The vaccine delivery sites (oral and injection) of all test and control animals were examined regularly after virus exposure for evidence of irritation or inflammation. The only finding of note was in one animal that had transient and mild mucosal changes consistent with blistering, observed on the upper gingiva (2 × 4 mm) day 11 post vaccination with complete resolution at the next exam at day 21, and ulceration (4 mm) of the buccal mucosa day 35 post vaccination which resolved by day 56 and did not alter feeding habits or interfere with daily activities. Importantly, there was no infectious virus recovered from the saliva or buccal swabs from this animal, nor from any body fluids collected from any of the experimental animals (serum, buccal, nasal, rectal swabs) at any time point. Complete blood counts remained within normal limits throughout follow-up post vaccine administration and the renal and hepatic functional measurements did not indicate any adverse reaction to immunization which was confirmed by histology. Overall, there was minimal vaccine toxicity with oral administration of the VSV-SARS2(+G) virus particles and the vaccine was well-tolerated in cynomolgus macaques.

Overall, our data support the clinical advancement of VSV-SARS2(+G) for oral booster vaccination of individuals previously infected or vaccinated but who have low or rapidly waning antibody titers. Previous studies have shown that peak post-SARS-CoV-2 infection and post-vaccination neutralizing antibody titers to SARS-CoV-2 differ between individuals based on age, infection severity and vaccine identity [18]. Antibody titers have also been shown to fall quite rapidly over time with half-lives varying from 2.5 and 6 months, signifying 4-fold to 25-fold titer reductions in the first year [4, 6, 19]. Thus, while the appropriate timing of SARS-CoV-2 vaccine booster shots has not yet been determined for specific subgroups of individuals, booster vaccination will likely be an important part of the strategy for longer-term control of the pandemic. Future studies comparing the efficacy of oral booster doses of VSV-SARS-CoV-2 and VSV-SARS2(G) in non-human primates would also be warranted after priming with currently approved vaccines.

Compared to currently approved mRNA and adenoviral vaccines which are administered by intramuscular injection, mucosal vaccination may offer a more reliable and durable defense against SARS-CoV-2 infection [3]. Mucosal surfaces in the nose and mouth provide the primary portal of entry for respiratory pathogens such as SARS-CoV-2 and are best protected by secretory polymeric IgA in mucus and saliva. Secretory polymeric IgA is efficiently induced by mucosal vaccines because they interact directly with mucosal associated lymphoid tissues such as the lingual and palatine tonsils which encircle the posterior outlet of the oral cavity. Oral mucosal vaccination may also overcome some of the logistical drawbacks of injectable vaccines such as the limited availability and high cost of vials, needles, syringes, and trained personnel, and the high prevalence of needle phobia in preadolescents, adolescents and college students. Comparing to intranasal vaccine delivery which has been successfully developed and deployed for influenza prevention [20], oral delivery may be a preferred approach to avoid the risks (and costs) of live virus aerosolization and CNS exposure associated with intranasal delivery.

Interestingly, while it was generally accepted at the outset of the SARS-CoV-2 pandemic that protein, mRNA and nonreplicating adenoviral vaccines could be developed very rapidly, it was also argued that vaccines built on replicating viral platforms might prove to be more potent, potentially affording protection after a single dose. Based on the extraordinary success of VSV-EBOV, a G-substituted VSV encoding the Ebola surface glycoprotein, which fully protects against Ebola virus infection within 2 weeks after a single intramuscular shot [21], it was predicted that a G-substituted VSV encoding the SARS-CoV-2 spike glycoprotein would be equally effective as an intramuscular vaccine to protect against COVID-19. However, in human trials the efficacy of this G-substituted construct, administered by intramuscular injection, fell short of expectations and induced suboptimal titers of neutralizing antibodies [22]. Our studies in cynomolgus macaques confirm the low immunogenicity of the VSV-SARS2 vaccine construct, but further extend the narrative by demonstrating the enormous promise of the same construct as an oral mucosal booster vaccine when modified to incorporate the VSV.G protein. We have therefore recently engineered a series of VSV-SARS2 constructs encoding the spike proteins of emerging SARS-CoV-2 variants. G protein pseudotyping of a replication defective VSV encoding the SARS-CoV spike glycoprotein was shown previously to enhance the immunogenicity of the construct when administered to NHPs by IM injection, but oral administration was not attempted [23]. Interestingly, another group recently exploited G protein trans-complementation as a way to restore the immunogenicity of a G-deleted VSV-derived vaccine for SARS-CoV-2 in ACE2 receptor negative mice [24]. VSV-eGFP-SARS-CoV-2, an autonomously replicating ACE2 receptor-dependent VSV which encodes the SARS2 spike glycoprotein and a GFP reporter gene was trans-complemented with “a small amount” of VSV-G protein, then used to vaccinate mice via the intraperitoneal route which resulted in robust, protective anti-spike antibody responses. These investigators further demonstrated the feasibility of intranasal vaccination using the same construct, but without G protein supplementation, in mice that were transgenic for the ACE2 receptor [24]. Although not formally proven, we speculate that the superior immunogenicity of the G-supplemented virus preparation, both by IM and PO routes, reflects a higher efficiency of muscle cell and oral mucosal transduction due the virus-displayed G protein interacting with the LDL receptor on myocytes and oral mucosal epithelium. We further speculate that the high potency of the intramuscular VSV-EBOV vaccine is likely the consequence of a high abundance of Ebola virus receptors in muscle tissue leading to efficient transduction by the injected viral particles, and that conversely the inferior efficacy of the VSV-SARS2 vaccine in the absence of G protein supplementation, is the consequence of low muscle expression of ACE2, the primary receptor for the SARS-CoV-2 spike protein [14, 25]. However, we cannot rule out the possibility of an additional contribution due to an immune adjuvant effect of the G protein itself.

The absence of an immune response to the VSV G protein when the G-supplemented viruses were administered orally is an important feature of the VSV-SARS2(+G) system which will allow for the future generation and deployment of G-supplemented VSVs whose G cistron has been substituted for envelope glycoproteins from other coronaviruses (e.g. those causing the common cold), or from pathogenic viruses belonging to non-coronaviridae families. G-supplemented VSV particles may therefore eventually prove to be a versatile oral mucosal vaccine platform capable of addressing a range of respiratory pathogens.

In summary, as a proof of concept, we utilized a relevant translational NHP model to demonstrate the safety and oral immunogenicity of a VSV-derived mucosal vaccine for SARS-CoV-2. Since completing these studies, we have developed a process for the scaled (GMP) manufacture of VSV-SARS-2(+G) in 293 suspension cultures and we have demonstrated product stability for up to 2 weeks at 4°C. Pre-IND discussions with FDA are ongoing.

## ACKNOWLEDGEMENTS

We gratefully acknowledge the excellent and expert contributions of Carlen Hill, Jody Janecek, Ruby Klish, Rachael Lee, Christian Moses, Brenna Mulhollam, Lucas Mutch, Melanie Niewinski, Sierra Palmer, Scott Oppler, Jordan Truell for husbandry and clinical care of our animals, Mikayla Chavis, Sarah Gresch, Laura Hocum Stone for in vitro studies, Meghan Moore and Timothy O’Brien for pathology, and Mellani Lubaug, Margret Tavai-Tuisalo’o, Jade Wilder for administrative support at the University of Minnesota’s Preclinical Research Center. We gratefully acknowledge the excellent and expert support of Parthasarathy Rangarajan, Naoya Sato, Anna Tran, Ravi Maisuria, Amar Singh, and Janice Anoka for sample processing and specimen archiving.

## AUTHOR CONTRIBUTIONS

SJR, KWP, CK, AB, LR, TC, RV, MG conceptualized and designed the experiments; JJ, RL, TC, RV, KS, CL, SReiter, RN, LS, TS, CP, PL, CG, CZ, MH, MT, JR, LZ, SW, BB, KK, SRamachandran, NP, MÁM-A performed research, TC, RV, KS, CL, KWP, SJR, MG analyzed data; SJR, KWP, TC, RV, NP, MÁM-A wrote the paper.

### Data and material availability

The IMMUNO-COV™ assay is available as a clinically validated test through a physician order.

## SUPPLEMENTARY MATERIALS

### Materials and Methods

#### Virus generation and characterization

Full-length human codon-optimized SARS-CoV-2 Spike (S) glycoprotein (NC_045512.2) in pUC57 was obtained from GenScript (MC_0101081). The plasmid was used as a PCR template to generate a cDNA encoding SARS-CoV-2 spike with a deletion in the nucleotides encoding the C-terminal 19 amino acids (S-Δ19CT) and 5’ MluI and 3’ NheI restriction sites. To generate the viral genome, the amplified PCR product containing S-Δ19CT was cloned into pVSV in place of VSV-G using the MluI and NheI restriction sites (Fig. 1). Plasmid was sequence verified and used for infectious virus rescue on BHK-21 cells as previously described [26]. VSV-G was co-transfected into the BHK-21 cells to facilitate rescue but was not present in subsequent passages of the virus. The viruses were amplified and propagated in Vero cells. The amplified viruses do not have VSV (G) glycoprotein and depend on SARS-CoV-2 spike (S) glycoprotein for entry and infection. To generate VSV-SARS2(+G), Vero producer cells were electroporated with a plasmid encoding VSV.G the day before infection with VSV-SARS2. Viruses were harvested 48h post infection and purified prior to use in experiments.

Correct incorporation of VSV G, N and M proteins and S glycoprotein in virions or virus infected cells was analyzed by Western blotting using a mouse α-SARS-CoV-2 Spike (1:1000, GeneTex #GTX632604), rabbit polyclonal α-VSV-G antibody (1:20,000, Abcam #ab83196), and rabbit polyclonal α-VSV antiserum (1:8000, Imanis #REA-005). Secondary antibodies were goat α-mouse IgG-HRP (1:30,000, Prometheus #20-304) and goat α-rabbit IgG-HRP (1:30,000, Prometheus #20-303). Membranes were developed for 2 min at room temperature using ProSignal^®^ Dura ECL Reagent (Prometheus #20-301). Protein bands were imaged using a BioRad ChemiDoc Imaging System. Flow cytometry was performed on infected cells at 8h post infection. Cells suspensions for flow cytometric analysis were prepared by lifting cells with Versene solution (Gibco #15040066) followed by staining with a monoclonal antibody against spike protein conjugated to DyLight™ 633.

Inhibition of virus infectivity on Vero cells was performed as follows. Vero-DSP1 and Vero-DSP2 co-monolayers, which produce functional Renilla luciferase following virus-mediated cell fusion, were seeded at 6×10^4^ cells per well in 96-well black-walled plates the day before assay. Neutralizing antibody cocktails were prepared in OptiMEM as follows: α-SARS-CoV-2 spike cocktail (mAb10914 at 6 µg/mL combined with mAb10922 at 2 µg/mL, Genscript) [4], α-VSV-G cocktail (mAbVSV-G 8G5F11, Absolute Antibody #Ab01401-2.3) at 2 µg/mL combined with mAb VSV-G IE9F9 (Absolute Antibody #Ab01402-2.0) at 10 µg/mL [27], and α-SARS-CoV-2 spike and α-VSV-G cocktail (combination of all four antibodies). Three-fold serial dilutions of the antibody cocktails were prepared and incubated with an equal volume of VSV-SARS2 or VSV-SARS2(+G). Final antibody concentrations in the virus mixtures were 3 µg/mL mAb10914, 1 µg/mL mAb10922 and mAbVSV-G 8G5F11, or 5 µg/mL mAbVSV0G IE9F9, and the final concentration of virus was 200 pfu/100 µL. Virus mixed with OptiMEM alone was used as a control. After a 30 min incubation at room temperature, 100 µL of each virus mix was overlaid onto the Vero-DSP1/2 co-monolayers in duplicate. Plates were returned to a 37°C/5% CO_2_ incubator. After 22 hours, EnduRen™ live cell substrate (6 mM, Promega #E6482) was added to wells, and luminescence was read after an additional 2 hours using a Tecan M Plex instrument (1000 ms integration, 100 ms settle time per well). Relative luciferase activity (infection) was determined for each condition relative to the virus in media only control.

#### Nonhuman primate study

##### Ethics statement

This study was conducted as approved by the University of Minnesota Institutional Animal Care and Use Committee. All animal experiments were executed in an Association for Assessment and Accreditation of Laboratory Animal Care–approved facility by qualified staff, following the guidelines and basic principles in the National Institutes of Health (NIH) Guide for the Care and Use of Laboratory Animals [28], the Animal Welfare Act, the U.S. Department of Agriculture, and the U.S. Public Health Service Policy on Humane Care and Use of Laboratory Animals. The University of Minnesota Institutional Biosafety Committee (IBC) approved work with VSV strains under BSL2 conditions and was performed according to IBC-approved standard operating procedures.

##### Animals

Twenty-two adult male and female adult Mauritian origin cynomolgus macaques (Macaca fascicularis) were allocated for this study. They were socially housed in same sex pairs or groups in climate-controlled rooms with a fixed light-dark cycle (12/12 hours) with a 30 minute sunrise illumination and sundown fade. Commercial monkey chow, 2055C Teklad Global Certified 25% Protein Primate Diet from Harlan Laboratories, and fruits or vegetables were provided twice daily and water was available ad libitum. The behavioral management program included social housing in pairs (except during bond breakdown or pending reintroduction), enrichment, and basic training. Environmental enrichment included human interaction and a variety of treats, toys, foraging puzzles, and sensory stimuli like music. Structural enrichment included perches, swings, and climbing apparatus. NHPs were trained prior to study for cooperation with routine husbandry tasks only which included targeting and shifting [29, 30].

In experiment 1, eighteen adult males and females with the median age of 3.7 years (range 3.4 to 6.7 years) and median body weight of 4.8 kg (3.2 to 6.8kg) were stratified by sex and randomly assigned to control and vaccine groups (Fig. 2). Animals received Sham control (N=6) at an equivalent volume to vaccinated animals, animals in vaccine groups received VSV-SARS2 at a dose of 10^7^ TCID_50_ PO (N=6) or 10^7^ TCID_50_ IM (N=6) on day 0. All 12 vaccinated animals received an oral dose of VSV-SARS2(+G) as an oral boost on day +42. In experiment 2, four demographically similar males with median age of 4.4 years (range 4.4 to 5.2 years) and median weight of 5.5kg (range 5.1 to 6.1 kg) were used (Fig. 3). Animals received VSV-SARS2(+G) at a dose of 10^8^ TCID_50_ PO (N=2) or IM (N=2) on day 0. Intramuscular vaccines were administered with a needle and syringe without adjuvant or orally using an oral syringe and gently tipping the head slightly back to deliver under the tongue and into the cheek pouches, with gently swabbing around the mouth with a cotton swab during the 5 minute dwell to mimic swishing.

##### Safety evaluation

Clinical observations of animals were performed at least twice daily for general attitude, activity level, gait, posture, appearance, feces, urine and any signs of pain or distress. Body weight was measured at least weekly. Animals were sedated for sham or test article administration, blood collection, and clinical exams using ketamine (5-15mg/kg IM) and midazolam was added (0.1-0.3mg/kg IM), if necessary, to extend sedation. All exams included weight and temperature measurements, complete examination of the oral mucosa, swabs, and blood sampling. Blood was collected using a peripheral vein and standard catheter at qualification, baseline prior to vaccination, then post vaccination biweekly, weekly, monthly or bi-monthly based on phase for clinical pathology that included complete blood counts, blood chemistries, CRP, and coagulation profiles. At these same timepoints nasal, buccal, and rectal swabs and fresh stool were also collected. At selected timepoints, additional assessment of immune profile pre and post vaccination was performed that included neutralizing antibody response and CD4/CD8 T cell response and characterization of VSV viremia and viral shedding.

##### Study Termination

All animals were euthanized at the completion of study and a full necropsy was performed which included a full gross examination. Bronchial alveolar lavage (BAL) was performed at necropsy by insertion of tube into the trachea, past the third bifurcation, and subsequent installation of 5-10 ml of sterile saline. Manual suction was applied to retrieve the BAL sample. Harvested tissues were fixed for a minimum of 72 hours in 10% neutral-buffered formalin and then embedded in paraffin. These tissues were evaluated histologically for signs of pathology by a board-certified veterinary pathologist blinded to study group allocations.

#### NHP Antigen Binding ELISA against SARS-CoV-2 Spike Trimer

Nunc Maxisorp microtiter plates (ThermoFisher Scientific) were coated with 100 ng/well of recombinant SARS-CoV-2 spike trimer protein (Cat.# SPN-C52H2, Acro Biosystems) in 200mM carbonate-bicarbonate buffer, pH 9.4. overnight at 4°C. Plates were washed and blocked with 1× SuperBlock™ in PBS for 30 minutes at room temperature (RT). Plates were washed again and incubated with serial dilutions of NHP sera or bronchiolar lavage fluid (BALF) diluted in sample buffer (1× SuperBlock™ T20 in PBS) and incubated for 90 minutes at RT. Plates were washed three times with PBS with 0.05% Tween 20 and then incubated for 45 minutes at RT with either horse radish peroxidase (HRP)-conjugated anti-NHP IgG (1:10,000, Cat.# PA1-84631, Invitrogen) or HRP-conjugated anti-NHP IgA (1:5,000, Cat.# 5220-0332, SeraCare) secondary antibodies diluted in sample buffer. After a final washing step, plates were developed using 100μl of SureBlue™ 1-Step TMB Substrate (3,3’,5,5’-tetramethylbenzidine; SeraCare) and the reaction stopped with an 100µL volume of HCl TMB Stop Solution (SeraCare) before the optical density (OD) was read at 450nm with a 630nm reference wavelength nm using an Infinite M200Pro microplate reader (Tecan). The dilution titers from serum IgG were determined using a cut point established for each animal based on Day 0 serum collection (collected prior to vaccination) and then multiplied by a cut point factor. For analysis of BALF IgA, the cut point was obtained using the average signal of the control group animals multiplied by a cut point factor. Cut point factors were determined separately for each antibody type and matrix using 95% confidence intervals obtained from individual analyses of the seronegative samples (pre-immune or saline control animals, respectively). Pre-immune serum paired to each animal was included on each analytical run to establish the plate-specific cut point on each assay plate. Dilution titers were calculated by fitting the data of all dilutions of a sample with a 4-parameter logistic curve to determine the dilution where absorbance signal falls below the cut point. Due to limited sample volume in evaluating IgA from the buccal swabs, the raw OD450 (with subtraction of 630nm reference) was compared directly between swab preparations prepared from saline control animals with swab preparations prepared from animals that received the oral VSV-SARS2(+G) boost.

#### IMMUNO-COV™ Neutralization Assays of NHP Samples

Vero-ACE2 cells were seeded at 10^4^ cells/well in 96-well black-walled plates with clear bottoms 16-24 h before being used for neutralization assays. On the day of assay, increasing dilutions (1:80, 1:160, 1:320, 1:640, 1:1280, and 1:2560) of serum samples were prepared and mixed with VSV-SARS2-Fluc in U-bottom suspension cell culture plates to a final volume of 240 µL/well. Virus, test samples, and controls were all diluted as appropriate in OptiMEM to generate final concentrations. Serum dilutions represent the final dilution following mix with virus. Virus was used at 300 pfu/assay well (300 pfu/100 µL in U-well mixtures). Pooled seronegative NHP serum was used as a negative control, and pooled seronegative NHP serum spiked with monoclonal anti-SARS-CoV-2 spike antibody was used as a positive control. Virus mixes in U-well plates were incubated at room temperature for 30-45 min, and then 100 µL of mixes were overlaid onto the Vero-ACE2 monolayers in duplicate. All time points for each animal were assayed on the same plate. Plates were returned to a 37°C/5% CO_2_ incubator for 24-28 hours. D-luciferin was then added to wells and luminescence was read immediately (30-90 seconds) using a Tecan M Plex or Tecan Lume instrument (100 ms integration, 100 ms settle time per well). Virus neutralizing titers were determined for each sample from an individual animal by calculating the dilution that resulted in luciferase signal greater than 50% of the Day 0 (prior to treatment) sample for that specific animal (IC_50_). The dilution was calculated by fitting all dilutions to a 4-parameter logistic curve fitting model.

#### VSV-G Antibody Neutralization Assays of NHP Samples

NHP serum samples and positive control rabbit α-VSV antiserum (Imanis #REA-005) were heat inactivated at 56°C for 30 min. Two-fold serial dilutions of samples and positive controls were prepared in quadruplicate starting at 1:10. Dilutions were mixed with equal volumes of VSV-GFP, such that the final VSV-GFP concentration was 400 TCID_50_/100 µL and the final serum and positive control concentrations were 1:20, 1:40, 1:80, 1:160, 1:320, 1:640, 1:1280, 1:2560, 1:5120, 1:10240, and 1:20480. Virus mixed with media alone was used as a negative control. After a 1 hour incubation at 37°C, 2×10^4^ BHK-21 cells were overlaid onto the virus mixes. Plates were incubated at 37°C/5% CO_2_ for 48 hours, at which time each well was scored for the presence or absence of cytopathic effects (CPE; cell death).

#### Infectious Virus Recovery (IVR) Assays

Buccal, rectal, and nasal swabs were cut and placed in 700 µL of OptiMEM containing antibiotics. After a 1 min. vortex at maximum speed, swabs were snap frozen in liquid nitrogen and then thawed at room temperature. Samples were centrifuged at 12,000 x g for 5 minutes at 4°C, then filtered through a 0.22 µm syringe filter. Testing of swab samples spiked with VSV-ΔG-SARS-CoV-2-S indicated that this method of virus extraction was compatible with recovery of infectious virus from swabs. Ten-fold serial dilutions starting at 10^−1^ of the samples were overlaid onto monolayers of Vero-αHis cells plated the day before. VSV-SARS-CoV-2-S-Δ19CT [17] was used as a control. Plates were incubated at 37°C/5% CO_2_ for 24 hours, at which time media was aspirated from all wells. Wells were washed with OptiMEM and then 100 µL of OptiMEM with 0.0004% trypsin/EDTA was added to each well. Plates were incubated for an additional 3 days at 37°C/5% CO_2_ before each well was scored for the presence of syncytia, indicative of infectious virus. Remaining preparations from buccal swabs prepared for IVR were used to evaluate NHP IgA against SARS-CoV-2 spike trimer in a binding ELISA assay.

#### Mice study

All animal procedures were reviewed and approved by the Mayo Clinic Institutional Animal Care and Use Committee. Female and male Ifnartm-CD46Ge mice [31] received intra-peritoneally 5×10^5^ TCID_50_ virus particles or 5μg of SARS-CoV-2 spike protein (Cat.# V0589-V08B, Sino Biological) adjuvanted with aluminum (Cat.#vac-alu-250,InvivoGen) on day 0 and 21. On days 21 (before boost) and 41, blood was collected. Parallel groups of mice were terminated on day 21 post-immunization for analysis of cellular immune response.

#### Mouse Antigen binding ELISA

Nunc ELISAs plates (ThermoFisher Scientific) were coated with 100 ng of recombinant SARS-CoV-2 spike protein (Cat.# V0589-V08B, Sino Biological) in 50mM carbonate-bicarbonate buffer, pH 9.6. overnight at 4°C. Plates were washed and blocked with 2% bovine serum albumin (BSA) in PBS for 2h at room temperature (RT). Plates were washed again and incubated with serial dilutions of mouse sera and incubated for 1h at 37°C. Plates were washed three times with PBS with 0.05% Tween 20 and then incubated for 1h at RT with horse radish peroxidase (HRP)-conjugated anti-mouse IgG (1:5,000, Cat.#62-6520, ThermoFisher Scientific), IgG1 (1:5,000, Cat.# 115-035-205, Jackson ImmunoResearch) or IgG2a (1:5,000, Cat.# 115-035-206, Jackson ImmunoResearch) secondary antibody. After final wash, plates were developed using 50μl of 1-Step Ultra TMB (3,3’,5,5’-tetramethylbenzidine; ThermoFisher Scientific) and the reaction stopped with an equal volume of 2M sulfuric acid before the optical density (OD) was read at 405 nm using an Infinite M200Pro microplate reader (Tecan). The endpoint titers of serum IgG responses were determined as the dilution that emitted an optimal density exceeding average of OD values plus three standard deviations of negative serum samples.

#### Pseudovirus production and neutralization assay

Pseudotyped lentiviruses with the SARS-CoV-2 spike protein were produced as described elsewhere [32]. In brief, HEK293T cells seeded overnight were transfected with HDM-Hgpm2, HDM-tat1b, pRC-CMV-Rev1b, pHAGE-CMV-Luc2-IRES-ZsGreen-W and a codon-optimized SARS-CoV-2 spike encoding the D614G amino acid change. The culture media was changed to fresh media containing antibiotics 18h to 24h post-transfection. At 60h post-transfection, supernatants were harvested and passed through a 0.45μm filter. For pseudovirus neutralization assay, three-fold serially diluted serum samples were mixed with SARS-CoV-2-pseudotyped-lentiviruses for 1 hours at 37 °C. And the mixture was added in quadruplicates to hACE-2-expressing HEK293T cells overnight, followed by media replenishment. At 60-72 hours-post incubation, the luciferase activity was detected by Bright-Glo Luciferase assay system (Cat.# E2610, Promega, Madison, WI, USA). The percentage of infection was calculated as the ratio of luciferase value with antibodies to that without antibodies. The half maximal inhibitory concentration (IC50) was determined by non-linear regression using GraphPad Prism version 8.4.2 for macOS (GraphPad Software, San Diego, CA, USA).

#### ELISpot Analysis

Frozen PBMCs were used to evaluate IFNγ, IL-4 T cell responses in NHP and frozen splenocytes were used IFNγ T cell responses in murine samples. Individual overlapping SARS CoV-2 Spike peptides (BEI Resources, NIAID, NIH: Peptide Array, SARS-Related Coronavirus 2 Spike (S) Glycoprotein, NR-52402; S1: 1 to 97 peptides, S2: 98 to 181 peptides), were pooled and dissolved in DMSO. Peptides were used at 0.5ug/ml concentration per peptide to elicit IFNγ and/or IL-4 responses. For NHP samples, low-fluorescent PVDF membrane (Millipore, Bedford, MA, USA) plates were coated with IFNγ and IL-4 capture antibodies overnight. Previously frozen PBMCs from NHP were washed, resuspended in T cell media (RPMI 1640 media containing 25 mM HEPES, NEAA, 2 mM L-glutamine, 1 mM sodium pyruvate, 0.05mM 2-mercaptoethanol supplemented with 10% (vol./vol.) heat-inactivated fetal bovine serum), counted and added to wells at 250,000 cells per well. SARS CoV2 Spike domain peptide pools (S1 and S2) were added to the plate and incubated at 37°C for 48 hours. Media containing equivalent DMSO and PMA/I were used as negative and positive controls respectively. Plates were then developed as per manufacturer’s instructions (MabTech Inc). Spot forming cells (SFC) were counted using automated MabTech IRIS reader with excitation/emission of 490nm/510nm for IFNγ and 550nm/570nm for IL-4.

For murine samples, ELISpot assays were performed using the mouse IFN-gamma ELISpot kit (Cat.# EL485, R&D systems Briefly, 5×10^5^ isolated splenocytes were added along with different stimuli in 200 μl of RPMI 1640 medium supplemented with 10% (vol./vol.) heat-inactivated fetal bovine serum for 48 h on each well of IFN-γ coated plates. Pooled 15-mer overlapping peptides from SARS-CoV-2 spike glycoprotein (BEI Resources) were used to stimulate splenocytes at 0.1μg of individual peptides/ml. As a positive control, PMA/Ionomycin cell stimulation cocktail (Biolegend, USA) was used at 2.5μl/ml, and as negative control, splenocytes were stimulated with equivalent DMSO concentration (0.8%). Post 48-hour incubation, plates were developed in accordance with manufacturer instructions. Developed IFN-γ spots were counted with an automated ELISPOT reader (CTL Analyzers LLC, USA). Each spot represented a single reactive IFN-γ–secreting T cell. Spots were normalized to DMSO control and figures represented as SFC per million cells for S1 and S2 peptide pools combined.

#### Statistical analyses

Tukey’s multiple comparison test or a two-tailed unpaired Student’s t test was conducted to compare differences between vaccine groups and the control group with Bonferroni correction applied to control the type I error rate for the comparisons.

## Supplemental Figure Legends

**Supplemental Figure 1:**
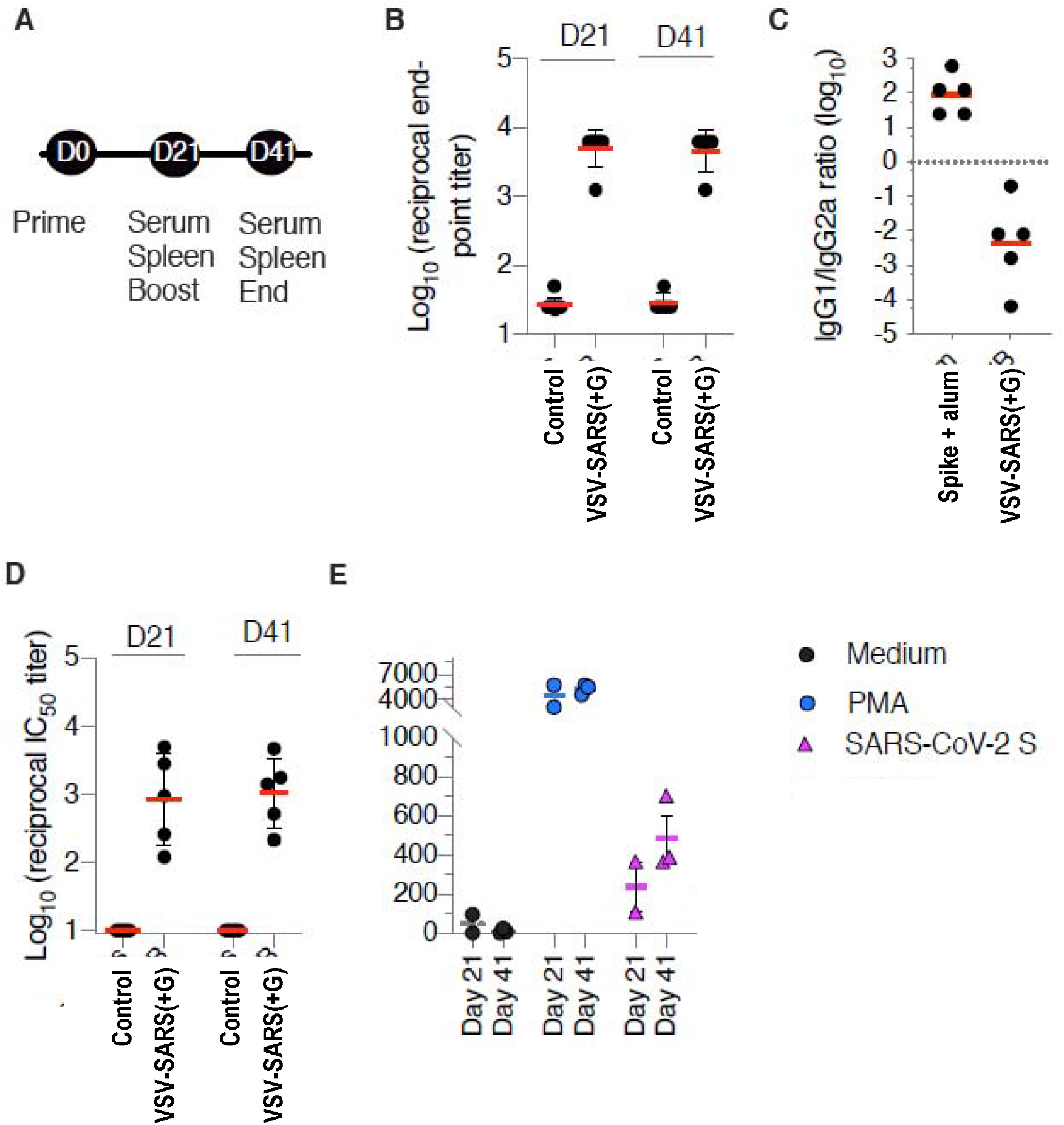
VSV-SARS2(+G) is powerfully immunogenic in mice. (**A**) Ifnartm-CD46Ge mice were vaccinated on day 0 and 21 with VSV-SARS2(+G) or irrelevant virus (control). Sera was collected on day 21 and 41. Spleens from vaccinated mice were collected from parallel groups. Collected sera were assessed by ELISA for SARS-CoV-2 spike-specific IgG (**B**), IgG1 and IgG2a (**C**). End-point titers (**B**) and end-point titer ratio of IgG1 to IgG2a (**C**) were calculated. Serum samples from animals immunized with SARS-CoV-2-Spike adjuvanted with Alum were used as control for a TH_2_ biased response. Pseudovirus neutralizing antibody responses were also calculated in vaccinated animals (**D**). In (**E**), splenocytes were isolated and stimulated with vehicle (media), PMA, or pools of overlapping peptides from SARS-CoV-2 spike. The number of cells expressing IFN--γ per 10^6^ splenocytes is displayed. Dots represent individual animals. Horizontal bars are mean per group.

**Supplemental Figure 2.**
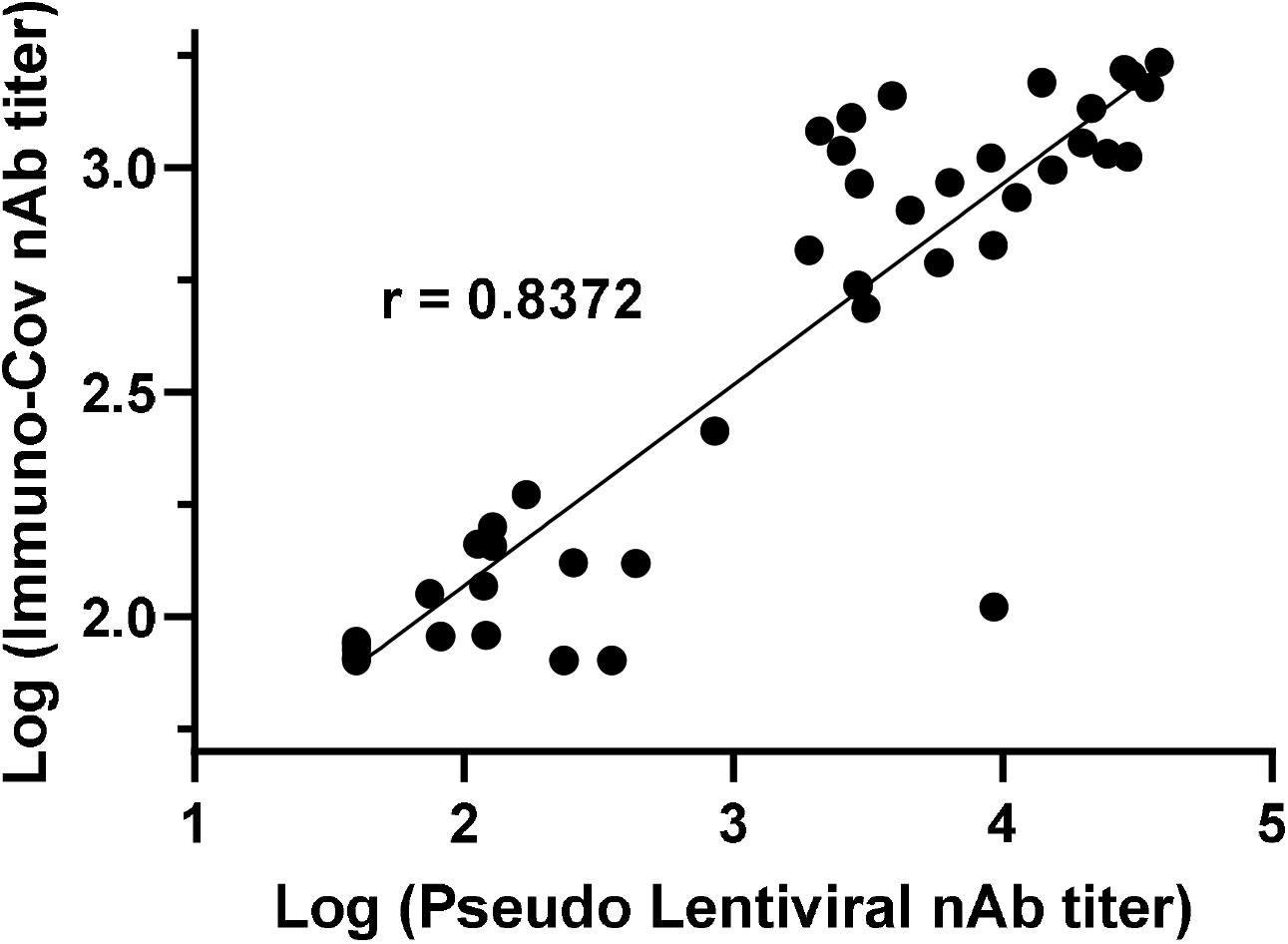
Good correlation in neutralizing antibody titers (nAb) measured using IMMUNO-COV assay that uses a VSV based versus a lentiviral based pseudovirus.

**Supplemental Figure 3.**
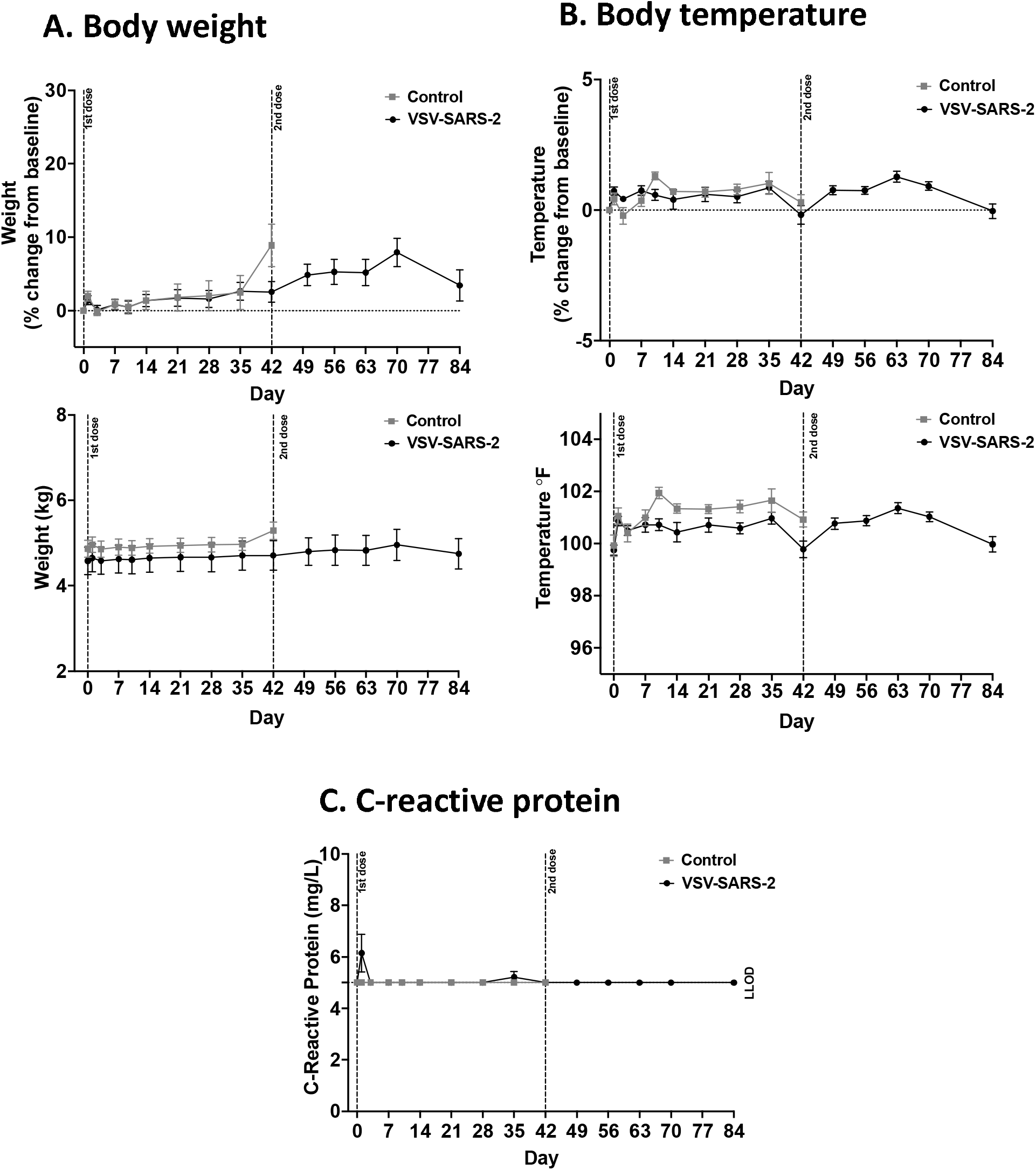
(**A**) Body weights, (**B**) temperature and (**C**) serum c-reactive protein in NHP after one prime dose of VSV-SARS(2) on day 0, and an oral boost of VSV-SARS2(+G) at day 42.

## REFERENCES

[1] Wouters OJ, Shadlen KC, Salcher-Konrad M, Pollard AJ, Larson HJ, Teerawattananon Y, et al. Challenges in ensuring global access to COVID-19 vaccines: production, affordability, allocation, and deployment. Lancet. 2021;397:1023–34.

[2] Ryan KA, Filipp SL, Gurka MJ, Zirulnik A, Thompson LA. Understanding influenza vaccine perspectives and hesitancy in university students to promote increased vaccine uptake. Heliyon. 2019;5:e02604.

[3] Mudgal R, Nehul S, Tomar S. Prospects for mucosal vaccine: shutting the door on SARS-CoV-2. Hum Vaccin Immunother. 2020;16:2921–31.

[4] Vandergaast R, Carey T, Reiter S, Lathrum C, Lech P, Gnanadurai C, et al. IMMUNO-COV™ v2.0: Development and Validation of a High-Throughput Clinical Assay for Measuring SARS-CoV-2-Neutralizing Antibody Titers. medRxiv. 2021:2021.02.16.21251653.

[5] Tan Y, Liu F, Xu X, Ling Y, Huang W, Zhu Z, et al. Durability of neutralizing antibodies and T-cell response post SARS-CoV-2 infection. Front Med. 2020;14:746–51.

[6] Widge AT, Rouphael NG, Jackson LA, Anderson EJ, Roberts PC, Makhene M, et al. Durability of Responses after SARS-CoV-2 mRNA-1273 Vaccination. N Engl J Med. 2021;384:80–2.

[7] Travis CR. As Plain as the Nose on Your Face: The Case for A Nasal (Mucosal) Route of Vaccine Administration for Covid-19 Disease Prevention. Front Immunol. 2020;11:591897.

[8] Durrant DM, Ghosh S, Klein RS. The Olfactory Bulb: An Immunosensory Effector Organ during Neurotropic Viral Infections. ACS Chem Neurosci. 2016;7:464–9.

[9] Baxter AL, Cohen LL, Burton M, Mohammed A, Lawson ML. The number of injected same-day preschool vaccines relates to preadolescent needle fear and HPV uptake. Vaccine. 2017;35:4213–9.

[10] Letchworth GJ, Rodriguez LL, Del cbarrera J. Vesicular stomatitis. Vet J. 1999;157:239–60.

[11] Roberts A, Buonocore L, Price R, Forman J, Rose JK. Attenuated vesicular stomatitis viruses as vaccine vectors. J Virol. 1999;73:3723–32.

[12] Finkelshtein D, Werman A, Novick D, Barak S, Rubinstein M. LDL receptor and its family members serve as the cellular receptors for vesicular stomatitis virus. Proc Natl Acad Sci U S A. 2013;110:7306–11.

[13] Fukushi S, Watanabe R, Taguchi F. Pseudotyped vesicular stomatitis virus for analysis of virus entry mediated by SARS coronavirus spike proteins. Methods Mol Biol. 2008;454:331–8.

[14] Hamming I, Timens W, Bulthuis ML, Lely AT, Navis G, van Goor H. Tissue distribution of ACE2 protein, the functional receptor for SARS coronavirus. A first step in understanding SARS pathogenesis. J Pathol. 2004;203:631–7.

[15] Yahalom-Ronen Y, Tamir H, Melamed S, Politi B, Shifman O, Achdout H, et al. A single dose of recombinant VSV-G-spike vaccine provides protection against SARS-CoV-2 challenge. Nat Commun. 2020;11:6402.

[16] Stevens TL, Bossie A, Sanders VM, Fernandez-Botran R, Coffman RL, Mosmann TR, et al. Regulation of antibody isotype secretion by subsets of antigen-specific helper T cells. Nature. 1988;334:255–8.

[17] Vandergaast R, Carey T, Reiter S, Lech P, Gnanadurai C, Tesfay M, et al. Development and validation of IMMUNO-COV: a high-throughput clinical assay for detecting antibodies that neutralize SARS-CoV-2. bioRxiv. 2020.

[18] Earle KA, Ambrosino DM, Fiore-Gartland A, Goldblatt D, Gilbert PB, Siber GR, et al. Evidence for antibody as a protective correlate for COVID-19 vaccines. medRxiv. 2021:2021.03.17.20200246.

[19] Lau EHY, Tsang OTY, Hui DSC, Kwan MYW, Chan WH, Chiu SS, et al. Neutralizing antibody titres in SARS-CoV-2 infections. Nat Commun. 2021;12:63.

[20] Carter NJ, Curran MP. Live attenuated influenza vaccine (FluMist(R); Fluenz): a review of its use in the prevention of seasonal influenza in children and adults. Drugs. 2011;71:1591–622.

[21] Suder E, Furuyama W, Feldmann H, Marzi A, de Wit E. The vesicular stomatitis virus-based Ebola virus vaccine: From concept to clinical trials. Hum Vaccin Immunother. 2018;14:2107–13.

[22] Herper M, Branswell H. In a major setback, Merck to stop developing its two Covid-19 vaccines and focus on therapies. 2021.

[23] Kapadia SU, Simon ID, Rose JK. SARS vaccine based on a replication-defective recombinant vesicular stomatitis virus is more potent than one based on a replication-competent vector. Virology. 2008;376:165–72.

[24] Case JB, Rothlauf PW, Chen RE, Kafai NM, Fox JM, Smith BK, et al. Replication-Competent Vesicular Stomatitis Virus Vaccine Vector Protects against SARS-CoV-2-Mediated Pathogenesis in Mice. Cell Host Microbe. 2020;28:465–74 e4.

[25] Zhou P, Yang X-L, Wang X-G, Hu B, Zhang L, Zhang W, et al. A pneumonia outbreak associated with a new coronavirus of probable bat origin. Nature. 2020;579:270–3.

[26] Ammayappan A, Nace R, Peng K-W, Russell SJ. Neuroattenuation of Vesicular Stomatitis Virus through Picornaviral Internal Ribosome Entry Sites. Journal of Virology. 2013;87:3217–28.

[27] Lefrancois L, Lyles DS. The interaction of antibody with the major surface glycoprotein of vesicular stomatitis virus. I. Analysis of neutralizing epitopes with monoclonal antibodies. Virology. 1982;121:157–67.

[28] I.f.L.A.R. Guide for the Care and Use of Laboratory Animals. Guide for the Care and Use of Laboratory Animals. Washington (DC) 2011.

[29] Graham ML. Positive Reinforcement Training and Research. In: Shapiro SJ, editor. Handbook of Primate Behavioral Management Boca Raton: CRC Press; 2017. p. 185–200.

[30] Graham ML, Rieke EF, Mutch LA, Zolondek EK, Faig AW, Dufour TA, et al. Successful implementation of cooperative handling eliminates the need for restraint in a complex non-human primate disease model. J Med Primatol. 2012;41:89–106.

[31] Mrkic B, Pavlovic J, Rulicke T, Volpe P, Buchholz CJ, Hourcade D, et al. Measles virus spread and pathogenesis in genetically modified mice. J Virol. 1998;72:7420–7.

[32] Crawford KHD, Eguia R, Dingens AS, Loes AN, Malone KD, Wolf CR, et al. Protocol and Reagents for Pseudotyping Lentiviral Particles with SARS-CoV-2 Spike Protein for Neutralization Assays. Viruses. 2020;12.

